# Experimental BTV-3 and BTV-8 infection of *Culicoides sonorensis* biting midges

**DOI:** 10.1101/2024.12.05.627042

**Authors:** Sophie Zeiske, Franziska Sick, Helge Kampen, Bernd Hoffmann, Martin Beer, Kerstin Wernike

## Abstract

BTV-3 emerged in the Netherlands in 2023 and spread rapidly to neighboring countries. Compared to the BTV-8 outbreak in 2006, the course of the BTV-3 epizootic is more severe. Experimental infection of laboratory-reared *Culicoides sonorensis* midges showed a slightly higher replication rate of BTV-3 than of BTV-8.

**One sentence summary line:** We experimentally infected laboratory-reared *Culicoides sonorensis* biting midges with bluetongue virus (BTV) serotypes 3 and 8 using virus-containing blood meal and found slightly higher replication rates of BTV-3.

## Main text

Bluetongue virus (BTV) is a non-contagious orbivirus that is transmitted between its mammalian hosts by *Culicoides* biting midges, causing severe disease in ruminant livestock (1). The first BTV outbreak ever recorded in central Europe in 2006 was caused by a serotype 8 strain (BTV-8) and led to a major epidemic (2). In 2023, a devastating BTV-3 outbreak started in the Netherlands and rapidly spread to neighboring countries (3). Arthropod-borne pathogens such as BTV are mainly spread by the dispersal of infected vectors and the movement of infected livestock (3). During vector monitoring in late 2023 in Germany near the Dutch border, BTV-3 was detected in a pool sample of *Culicoides* biting midges (4).

Comparison of the spread between farms of BTV-8 in 2006/2007 and BTV-3 in 2023 by transmission kernel analysis, which describes the distance-dependent probability of disease transmission from an infected farm to a susceptible farm, revealed a very similar kernel shape parameter of the BTV-8 and the BTV-3 outbreaks. This suggests that the mechanisms of disease spread through short distance dispersal of infected midges and other modes for longer distances, such as livestock movement, were similar between the two outbreaks (3). However, a much higher amplitude parameter was observed for the 2023 BTV-3 epidemic, indicating a faster disease spread. This could be due to higher temperatures of about 2°C above normal during the observed period (September-November) of the 2023 BTV-3 outbreak compared to the corresponding period of the 2006 BTV-8 outbreak. Another reason could be a higher infection and transmission efficiency of the midges for BTV-3, which would also result in a faster spread of the disease (3).

Since laboratory colonies of European biting midge vector species are not available, several experimental infection studies have been conducted using field-collected midges to investigate the vector competence of *Culicoides* species with different BTV-8 strains (5, 6). The laboratory-reared colony of *Culicoides sonorensis* is a suitable model to study infection dynamics under standardised laboratory conditions (7, 8), since this species plays a crucial role for BTV transmission in North America (9). Although most experimental BTV-8 infection studies using field-captured midges aim to calculate replication rates, the results of different studies are not easily comparable due to differences in experimental design, sample processing and data analysis. No studies are available for the more recent BTV-3 strain.

To directly compare the replication properties of BTV-8 and BTV-3 in biting midges, we performed infection experiments with the laboratory colony of *C. sonorensis*.

## The Study

A laboratory colony of *C. sonorensis* was reared in the BSL2 insectary of the Friedrich-Loeffler-Institut (FLI), Greifswald-Insel Riems, as described previously (10). Three-day-old biting midges were offered caprine (trial one) or ovine (trial two) heparin blood, obtained from the FLI, mixed 1:1 with BTV-8 or BTV-3 in cell culture medium (Minimum Essential Medium). Virus stocks of BTV-8 (strain BH311/06, isolated from a German sheep during the 2006 outbreak) and BTV-3 (4) were propagated on BHK-21 cells (RIE164, Collection of Cell Lines in Veterinary Medicine (CCLV), Friedrich-Loeffler-Institut, Greifswald-Insel Riems, Germany). The blood meal contained 10^6^ 50 % tissue culture infective dose per ml (TCID_50_/ml), which was confirmed by back-titration after feeding. As a negative control (NC), blood was mixed with virus-free cell culture medium. After pre-heating to 37°C, the blood meal was offered to the midges using a “Hemotek membrane feeding system” (Hemotek, Blackburn, UK) for 30 minutes. Midges were sorted under short-term CO_2_-anaesthesia on a cooling plate. Clearly engorged females were transferred to a new cage and kept for the course of the experiment. 16 blood-fed midges per group (BTV-8, BTV-3, NC) were processed immediately after feeding as uptake controls (Figure 1). Midges were kept inside gaze covered cages in an incubator at 27°C and a relative humidity of 85% with an 8h dark/ 16h light regime and supplied with 5% glucose ad libitum. After an incubation period of 6 days, surviving midges were harvested. All midges were placed individually in tubes containing 200 µl phosphate-buffered saline (PBS) and a 5 mm stainless steel ball (Figure 1). Following homogenization using a TissueLyzer (Qiagen, Hilden, Germany) for three minutes at 30 Hz, total RNA was extracted for each sample using the King Fisher 96 Flex (Thermo Scientific, Braunschweig, Germany) in combination with the NucleoMag VET kit (Macherey Nagel, Düren, Germany) according to the manufacturer’s instructions. The RNA extracts were analyzed by a BTV specific RT-qPCR (11) with an external full virus BTV-3 standard, which was used to calculate the number of BTV genome copies per midge.

**Figure 1.**
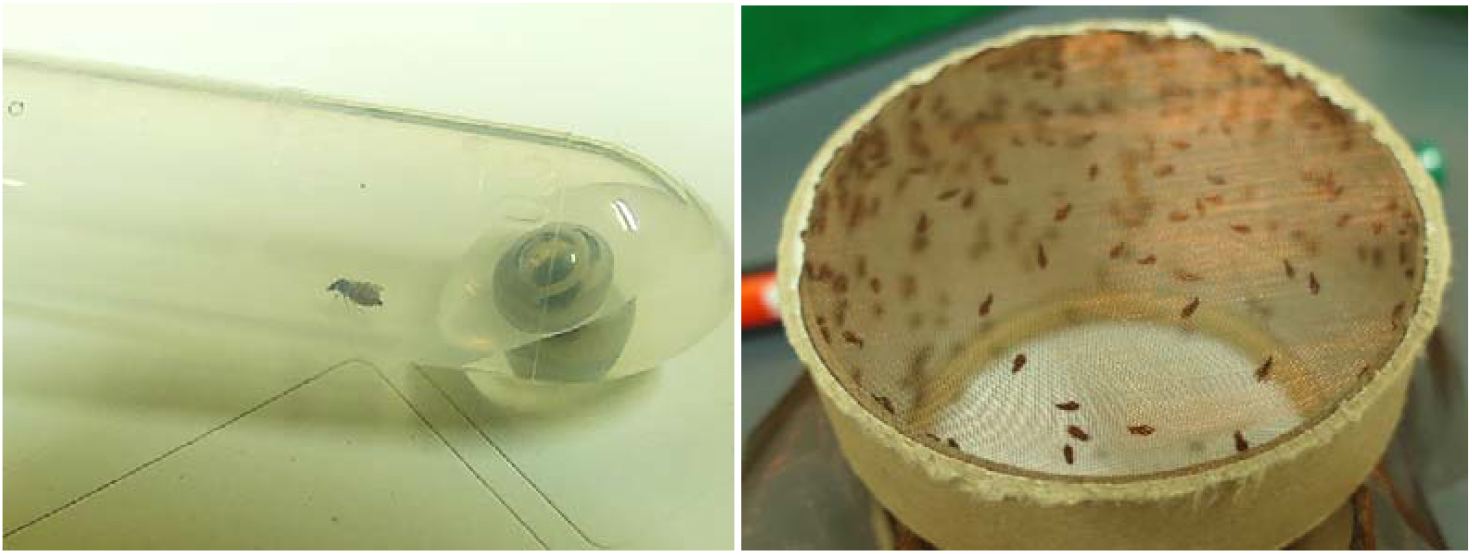
*C. sonorensis* after blood meal (left). The biting midge was placed individually in a tube containing 200 µl phosphate-buffered saline (PBS) and a 5 mm stainless steel bead for further processing. Engorged females in new netted cardboard cage after blood feeding (right).

Ingestion of virus-spiked blood (BTV-8 and BTV-3) led to PCR positivity in all midges harvested directly after the blood meal (day 0). The highest BTV genome copy number in an individual midge from a day-0-group was used as a cut off value to evaluate virus replication in midges fed with the same blood meal but harvested only after 6 days of incubation. Two consecutive trials with the same set-up were performed as biological replicates and to achieve a higher number of analyzable midges. In the first trial using BTV-3, 89 out of 319 surviving midges tested positive by RT-qPCR at 6 dpi, and 17 of these (5.32%) had viral loads higher than the day-0-group, indicating efficient virus replication. In the BTV-8 group of the first trial, 197 out of 330 surviving midges tested positive by RT-qPCR, and 12 of them (3.64%) showed efficient virus replication. In the second trial, in the BTV-3 group, 133 out of 250 surviving midges tested positive by RT-qPCR, and 8 of them (3.20%) showed virus replication. In the BTV-8 group of the second trial, 110 out of 222 surviving midges tested positive by RT-qPCR, and 7 (3.15%) of them showed virus replication. Midges of the negative control group tested negative by RT-qPCR at all times (Figure 2). Overall, 4.39 % of BTV-3 infected midges replicated the virus, while 3.44% of the BTV-8 infected midges replicated the virus (Figure 2). Statistical analysis with a two-sided Fisher exact test showed that the differences in the replication properties between BTV-3 and BTV-8 are not statistically significant.

**Figure 2.**
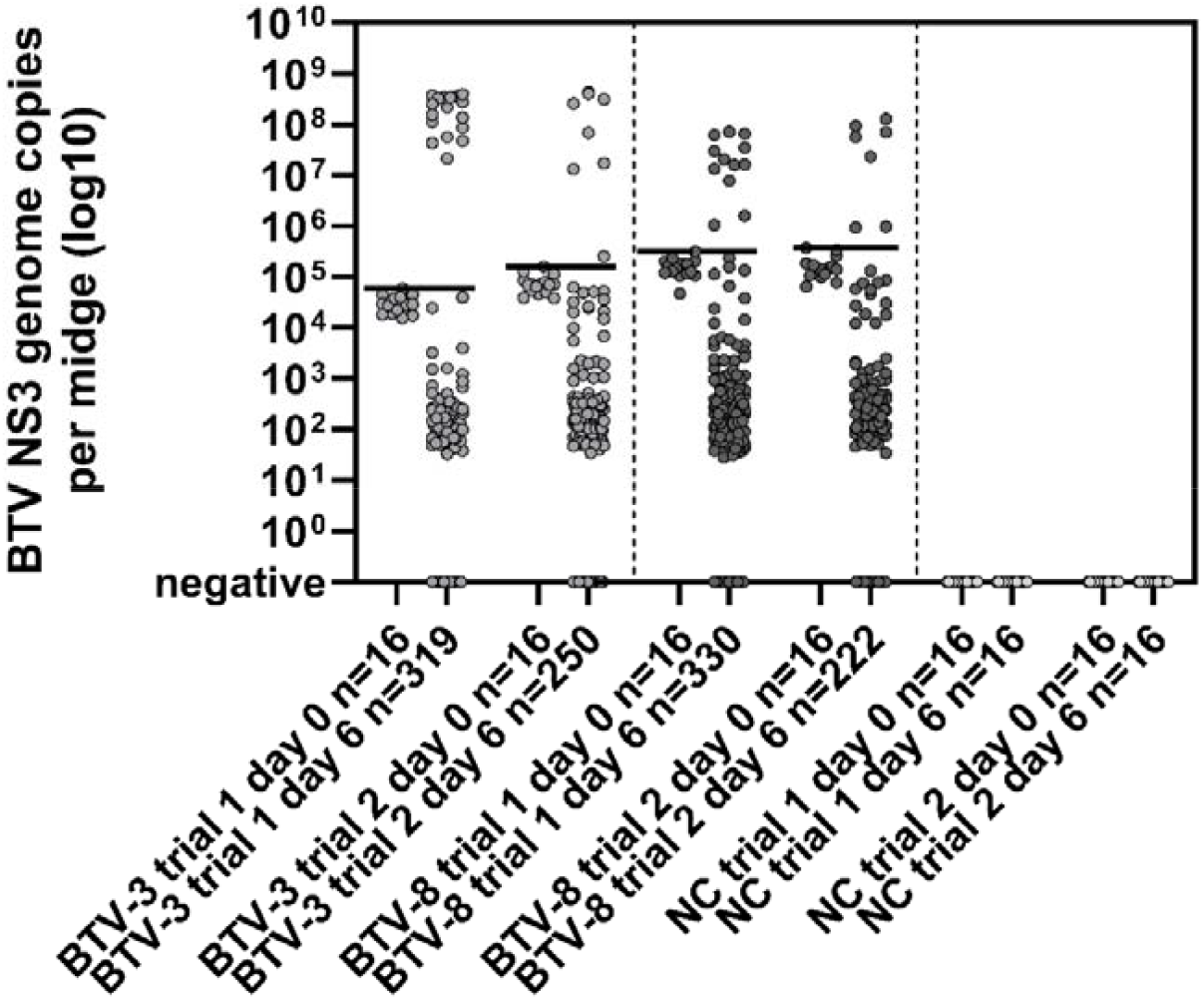
RT-qPCR results of midges experimentally infected with BTV-3 or BTV-8 spiked blood meal and midges fed with virus-free blood (negative control, NC). Individual midges were tested for BTV genome immediately after ingestion of the blood meal (day 0) or six days after the blood meal (day 6). Horizontal black lines indicate the highest BTV copy number measured in any of the midges of the respective group immediately after blood meal ingestion. The experiment was performed in two subsequent trials, the trial number is given in the label of the x-axis.

## Conclusions

In our study, orally infected midges were processed individually to calculate replication rates of BTV-3 and BTV-8, based on PCR-determination of genome copies per midge. In our laboratory colony of *C. sonorensis*, oral BTV-3 infection resulted in a slightly higher percentage of virus-positive biting midges with demonstrated replication than BTV-8 infection, with a total of 4.39% of BTV-3 infected midges and 3.44% of BTV-8 infected midges replicating the virus. The higher proportion of midges with BTV-3 replication may be a factor contributing to the observed faster outbreak progression of the current BTV-3 outbreak in comparison to the BTV-8 outbreak in 2006/2007.

## Acknowledgments

We would like to thank Uday Gottam and Ulrike Neumann for their excellent technical assistance. *Culicoides sonorensis* were originally developed and supplied by The Pirbright Institute under BBSRC project code: BBS/E/I/00007039. The study was funded by the German Federal Ministry of Food and Agriculture (BMEL) through the Federal Office for Agriculture and Food (BLE), grant number 28N207601.

